# SimpleForest - a comprehensive tool for 3d reconstruction of trees from forest plot point clouds

**DOI:** 10.1101/2021.07.29.454344

**Authors:** Hackenberg Jan, Calders Kim, Miro Demol, Pasi Raumonen, Alexandre Piboule, Disney Mathias

## Abstract

The here-on presented SimpleForest is written in C++ and published under GPL v3. As input data SimpleForest utilizes forestry scenes recorded as terrestrial laser scan clouds. SimpleForest provides a fully automated pipeline to model the ground as a digital terrain model, then segment the vegetation and finally build quantitative structure models of trees (QSMs) consisting of up to thousands of topologically ordered cylinders. These QSMs allow us to calculate traditional forestry metrics such as diameter at breast height, but also volume and other structural metrics that are hard to measure in the field. Our volume evaluation on three data sets with destructive volumes show high prediction qualities with concordance correlation coefficient *CCC* 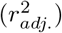 of 0.91 (0.87), 0.94 (0.92) and 0.97 (0.93) for each data set respectively.

We combine two common assumptions in plant modeling “*The sum of cross sectional areas after a branch junction equals the one before the branch junction*” (***Pipe Model Theory***) and “*Twigs are self-similar*” (***West, Brown and Enquist model***). As even sized twigs correspond to even sized cross sectional areas for twigs we define the *Reverse Pipe Radius Branchorder* (*RPRB*) as the square root of the number of supported twigs. The prediction model *radius* = *B*_0_ * *RPRB* relies only on correct topological information and can be used to detect and correct overestimated cylinders. In QSM building the necessity to handle overestimated cylinders is well known. The RPRB correction performs better with a *CCC* 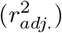 of 0.97 (0.93) than former published ones 0.80 (0.88) and 0.86 (0.85) in our validation.

We encourage forest ecologists to analyze output parameters such as the *GrowthVolume* published in earlier works, but also other parameters such as the *GrowthLength, VesselVolume* and *RPRB* which we define in this manuscript.

**Upload statement:** Self-uploaded pre-print for peer-review submitted manuscript. The manuscript was submitted on 26th of July 2021 to Plos Computational Biology:

I, Jan Hackenberg uploaded this manuscript because the automated journal upload was rejected for the following reason:

*Thank you for considering posting your manuscript “SimpleForest - a comprehensive tool for 3d reconstruction of tree from forest plot point clouds.” as a preprint. Your manuscript does not meet bioRxiv’s criteria and therefore we will not be sending it for posting as a preprint. For more information about our checks, see link*.

*We have noted that it contains material that is potentially subject to copyright. In particular, screenshot in Figure 1. Preprints posted to bioRxiv following submission to PLOS journals are done so under the CC BY license. To avoid a potential breach of the copyright that applies to the material listed above, we are unable to make the manuscript publicly available*.

Fig 1.
Submission system screenshot.

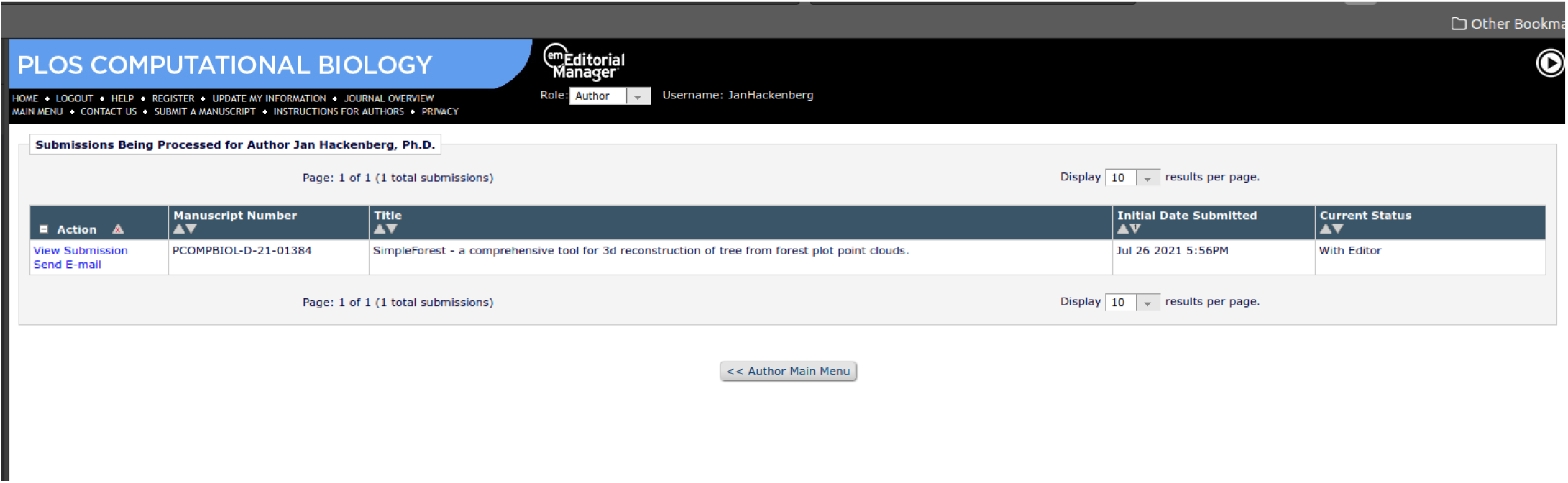

*Please note that this decision does not affect the editorial process at PLOS Computational Biology. Your manuscript is being separately assessed with regards to sending for peer review*.

From section Abstract on, the pdf you see is same as submitted one.

## 1 Introduction

Forests are key part of the global carbon cycle [1]. The quantification of global carbon bound in forests generally relies on forest inventory data. In traditional forestry often only two parameters per sampled tree are derived, the diameter at breast height (DBH) and the height (h). Above-ground biomass (AGB) is then derived either by a volume table look-up of DBH and h or by an allometric model utilizing the two parameters as input variables.

Such tables and formulas can be found published [2, 3] but this is not always the case. Especially when species provenance and the variation of regional growth patterns are taken into account, one has to rely on destructive data collection [4]. In the last two decades utilizing the point cloud data of terrestrial laser scans to derive directly the volume of trees has been shown as a fast, less costly and non destructive alternative [5].

The complete trees above-ground volume (AGV) can be directly assessed from the topological ordered cylinder structure of so called quantitative structural models (QSM)s [6, 7]. QSMs are not only important to collect AGV data which can be multiplied with density predictions [8] for AGB estimates, they allow forest ecology relevant insight into tree architecture itself [9, 10]. From a single TLS forest plot scan multiple thousands or even up to millions of geometrical measurements can be derived. Nevertheless, by purely using limited destructive measured data theories such as the West Brown and Enquist (WBE) model [11–13] or the pipe model theory [14, 15] have been developed under named limitations.

With QSMs new advances such as the rework of the WBE [16] describing a generalization for asymmetric branched trees are possible, to help understanding the tree architecture [9].

As an official successor of *SimpleTree* [7] we provide here the *SimpleForest* plugin for the *Computree* platform [17] to retrieve QSMs from plot level raw data with a fully automated pipeline consisting of various filters, a digital terrain model (DTM) algorithm, vegetation segmentation and finally two different QSM modelling approaches. Firstly, the so called SphereFollowing method [4,18] and additionally the Dijkstra [19] based method such as proposed by [20,21]. Our implementation of the latter one is known to perform worse than the first and is here-on undiscussed.

*SimpleForest*’s fully automatic processing chain can be inspected step by step with the powerful visualizer of *Computree. SimpleForest*’s QSMs can be conveniently extracted in csv format to be postprocessed with AMAPstudio [22] or R software [23]. Ply format allows visual inspection with tools like *CloudCompare, Meshlab* or *Blender*. The latter approach is useful for exemplary quality assessment after computation when a larger number of plots is processed in the batch-mode of *Computree*. The SphereFollowing parameters are automatically searched for in higher dimensional space. This automatic search is especially user friendly for users without informatic knowledge but also did outperform us on expert level.

In case wood vs. leaf separation is desired, separated tree clouds can be processed in a supervised machine learning manner. Remaining leaf fragments in the cloud lead to errors in QSMs which are afterwards automatically handled by filters relying on the WBE [24] theory. Here we use new invented parameters which show a better *Radius* correlation than traditional ones for the important diameter relations in scaling theories.

## Supporting information

### S1 Text. The SimpleForest UserGuide

- Over 100 pages of documentation are contained in the User Guide: https://gitlab.com/SimpleForest/computree/-/blob/master/bin/SimpleForestUserGuide.pdf, https://doi.org/10.5281/zenodo.5138255.

All algorithms are described in detail with a special focus on the parameterization of the steps.

#### 1.1 Data

We will evaluate on 5 different data sets. Each data set is online available, in one repository: https://doi.org/10.5281/zenodo.5131717.

##### S1 Dataset Hackenberg *et al*. 2015a data [35]

12 *Quercus petraea* (leaf-off), 12 *Erythrophleum fordii* (leaf-on) and 12 *Pinus massioniana* (needle-on) tree scans are available with harvested ground truth volume data [35]. The *Quercus petraea* trees have been overgrown with moss leading to larger overestimation of the *Radius*.

- *Erythrophleum fordii* clouds raw: https://zenodo.org/record/5131717/files/hackenbergErythrophleumRawClouds.zip
- *Erythrophleum fordii* clouds denoised: https://zenodo.org/record/5131717/files/hackenbergErythrophleumDenoisedCloudsCC.zip
- *Pinus massioniana* clouds raw: https://zenodo.org/record/5131717/files/hackenbergErythrophleumRawClouds.zip
- *Pinus massioniana* clouds denoised: https://zenodo.org/record/5131717/files/hackenbergPinusDenoisedClouds.zip
- *Quercus petraea* clouds denoised: https://zenodo.org/record/5131717/files/hackenbergQuercusDenoisedClouds.zip

##### S2 Dataset De Tanago *et al*. 2017 data [36]

29 tropical buttresses trees have been scanned across sites in Peru, Indonesia and Guyana. Harvested reference data is included, [36]. All trees have been scanned in leaf-on condition.

- Peru, Indonesia and Guyana clouds raw: https://zenodo.org/record/5131717/files/deTanagoRawClouds.zip
- Peru, Indonesia and Guyana clouds denoised: https://zenodo.org/record/5131717/files/deTanagoDenoisingScriptsDenoisedClouds.zip

##### S3 Dataset Demol *et al*. 2021 data [37]

65 trees in total of the species *Fraxinus excelsior*, *Fagus sylvatica*, *Pinus sylvestris* and *Larix decidua* have been scanned and published with reference data [37]. The *Pinus sylvestris* trees have been scanned in needle-on condition, the other species were scanned without foliage.

- Clouds: https://zenodo.org/record/5131717/files/demolClouds.zip

##### S4 Dataset Disney *et al*. 2018 data [38,39]

An *Acer pseudoplatanus* [39] tree cloud of a large and complex individual with DBH larger 1 m is used to show statistical analysis potential. The tree is located in Wytham Woods, UK.

- Clouds: https://zenodo.org/record/5131717/files/wythamCloud.zip

##### S5 Dataset High resoluted leaf-less plot data

A full plot in leaf off conditions. The plot does not have ground truth data in attached, but the quality of tree segmentation as well as DTM and QSM modeling can be visually assessed.

- Clouds: https://zenodo.org/record/5131717/files/leibzigCloud.zip

###### S1 Processing scripts SimpleForest scripts to process S1 Dataset

- *Erythrophleum fordii* denoising scripts: https://zenodo.org/record/5131717/files/hackenbergErythrophleumDenoisingScripts.zip
- *Pinus massoniana* denoising scripts: https://zenodo.org/record/5131717/files/hackenbergPinusDenoisingScripts.zip
- QSM modeling script: https://zenodo.org/record/5131717/files/hackenbergQsm.xsct2

###### S2 Processing scripts SimpleForest scripts to process S2 Dataset

- Denoising scripts: https://zenodo.org/record/5131717/files/deTanagoDenoisingScriptsDenoisedClouds.zip
- Poisson reconstruction buttress script: https://zenodo.org/record/5131717/files/deTanagoButtressPoisson.xsct2
- QSM modeling script: https://zenodo.org/record/5131717/files/deTanagoQsm.xsct2

###### S3 Processing scripts SimpleForest scripts to process S3 Dataset

- QSM modeling script: https://zenodo.org/record/5131717/files/demolQsm.xsct2

###### S4 Processing scripts SimpleForest scripts to process S4 Dataset

- QSM modeling script: https://zenodo.org/record/5131717/files/wythamQsm.xsct2

###### S5 Processing scripts SimpleForest scripts to process S5 Dataset

- QSM modeling script: https://zenodo.org/record/5131717/files/leibzigQsm.xsct2

###### S1 Validation scripts SimpleForest scripts to validate results of S1 Processing scripts

- Volume validation script: https://zenodo.org/record/5131717/files/hackenbergValidation.R

###### S2 Validation scripts SimpleForest scripts to validate results of S2 Processing scripts

- Volume validation script: https://zenodo.org/record/5131717/files/deTanagoValidation.R

###### S3 Validation scripts SimpleForest scripts to validate results of S3 Processing scripts

- Volume validation script: https://zenodo.org/record/5131717/files/demolValidation.R

###### S4 Validation scripts SimpleForest scripts to validate results of S4 Processing scripts

- Statistical plotting script: https://zenodo.org/record/5131717/files/wythamAnalysis.R

###### S5 Validation scripts SimpleForest scripts to validate results of S1 Processing scripts, S2 Processing scripts, S3 Processing scripts

- Combined results data table: https://zenodo.org/record/5131717/files/tableAll.csv
- Volume validation: https://zenodo.org/record/5131717/files/ValidationScriptAll.R

#### 1.2 Software

##### S1 Software Software code repository

- Under the GPL version 3 license: https://gitlab.com/SimpleForest/computree/-/blob/master/pluginSimpleForest/GPL_v3_template
- we provide source code with compilation instructions for the here presented *SimpleForestv5.3.1* plugin published: https://gitlab.com/SimpleForest/computree/-/commits/v5.3.1.
- Inside a subfolder this repository contains a Win10 compiled executable: https://gitlab.com/SimpleForest/computree/-/tree/master/bin.
- Persistent 5.1.3: https://doi.org/10.5281/zenodo.5138255

##### S2 Software Online resources for S1 Software

- An homepage: https://www.simpleforest.org/.
- A video tutorial channel: https://www.youtube.com/channel/UCq2dqxUF3IGmusX0ptk7xhw.
- Support forum *Computree:* http://rdinnovation.onf.fr/projects/computree/boards.
- *SimpleForest* users help users list: https://lists.posteo.de/listinfo/simpleforest.

## 2 Design and Implementation

*Computree* and *SimpleForest* rely on the following third party libraries: the Point Cloud Library (PCL) [25], the Boost library, the Gnu Scientific Library (GSL) [26], the OpenCV library, the QT library and the Eigen library.

All steps are implemented multi-threaded with an optimized asymptotic runtime to execute as fast as possible. In S1 Text we describe more than 20 steps regarding the IO data, with the underlying algorithms explained in detail and the parameterization is discussed as well. Here only a short overview is given.

### 2.1 Computree

*Computree* is a collaborative platform published under LGPL for 3d data processing for forestry applications. *Computree* can visualize multiple results of intermediate computation steps at once, Fig 2. If the user wants, he can display one result into a new visualizer windows each. As all visualizers can have a shared camera view the visual inspection is user-friendly implemented.

**Fig 2.**
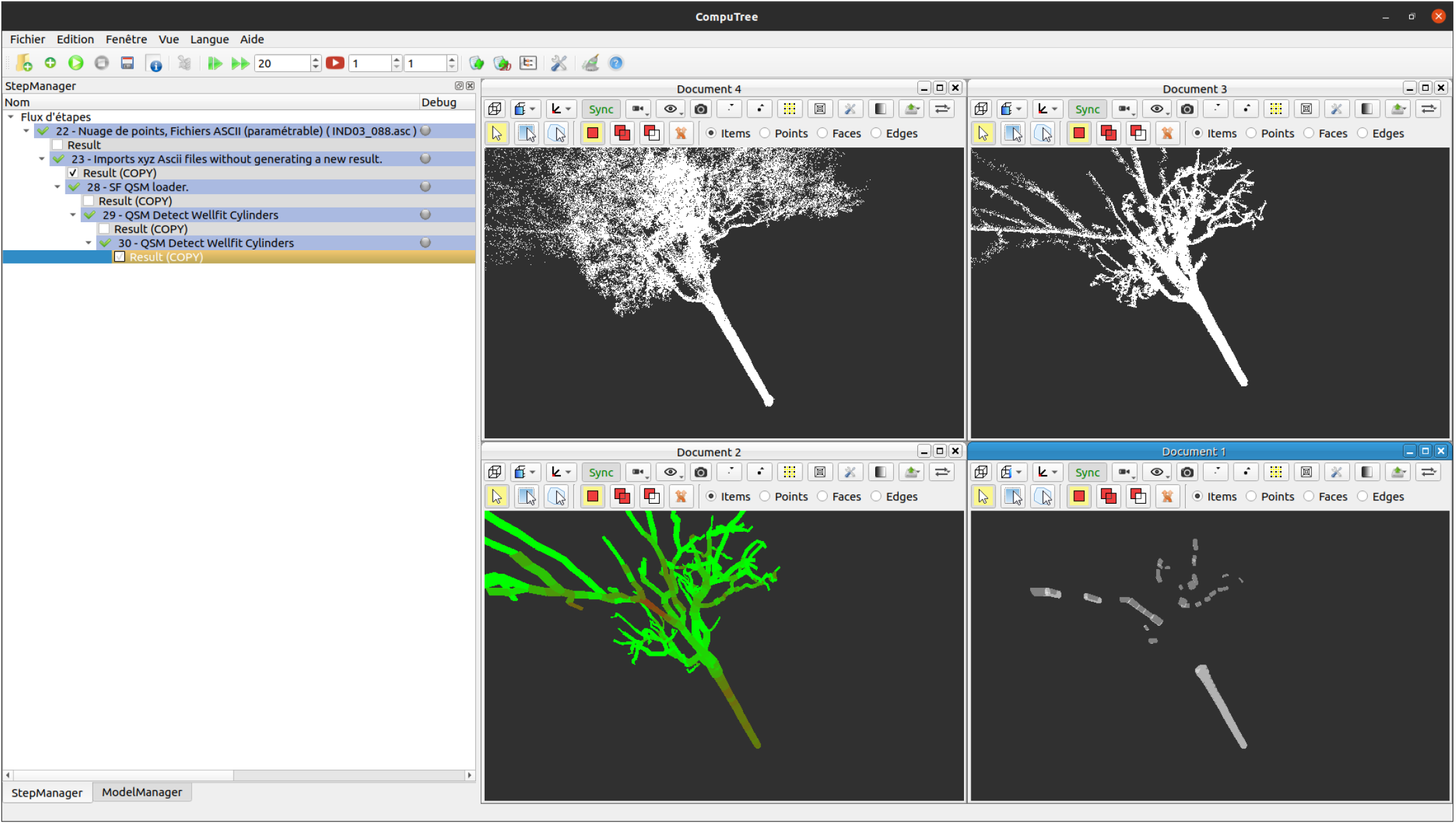
The user interface of *Computree*. Four visualizers with a shared camera view are opened. UpperLeft shows the leaf-on condition input cloud, UpperRight the de-noised cloud. On the LowerLeft we see the resulting QSM, with error colors. Brown means good fitted cylinders and green bad fitted ones. On the LowerRight the cylinders are depicted which are considered good enough for an allometric fit.

With the QT user interface steps parameters can be conveniently adjusted. Parameterized step pipelines can be saved and reloaded.

### 2.2 Point Cloud Filters

SimpleForest implements various point cloud filters for generating slices in a desired height above the DTM, filters to retrieve either ground or stem points by the normal deviation from the z-axis as well as frontends to few PCL [25] filters and the Belton filter [28].

#### 2.2.1 Gaussian Mixture Models with Fast Point Feature Histograms (FPFH) [27]

A semi automatic supervised machine learning technique utilizing gaussian mixture models [28], using the FPFH feature space and relying on a single range parameter. The filter was implemented for de-leaving. Up to ten automatic generated clusters have to be labeled by the user into LEAF, WOOD and UNCLASSIFIED. Then on each original point KNN clustering is performed, if more WOOD than LEAF points are in the neighborhood, the final classification is WOOD, otherwise LEAF.

### 2.3 DTM

A DTM can be computed via a pyramidal approach where raster cell size is halved iteratively from full plot size to a user desired threshold. Into points within one cell MLSAC planes are fitted [29]. To be accepted the plane has to surpass a closeness test to the according lower resoluted cell’s plane from the previous iteration.

### 2.4 Tree Segmentation

The segmentation consists of three main procedures, each implemented in at least a single step to be able to show intermediate results.

#### 2.4.1 Generating the seeds

We use as input a slice of points above the DTM at DBH. The slice points have been filtered with the stem point filter to remove undergrowth branch connections between neighboring trees. By euclidean clustering retrieved clusters serve now as seeds and each cluster receives an unique ID.

#### 2.4.2 Growing seeds to trees with Dijkstra [19]

On all points of the full vegetation Dijkstra’s [19]’s algorithm is executed in a competitive manner with the seed clusters of the previous step. A point receives the cluster ID of the closest seed point. The distance measure here is the length of the Dijkstra path to a connected seed point. To be able to jump over occlusion gaps the user can scale the cloud by z-coordinate. If the z-coordinate is multiplied with a factor smaller than one, vertical gaps have less impact compared to horizontal gaps. We want to achieve this, because natural non occlusion gaps are large in z-dimension for multi layered forests, but can be rather small in the x,y plane for neighboring trees.

#### 2.4.3 Classifying remaining points with Voronoi regions

Clusters too far away from the main vegetation cluster and not reached by the Dijkstra routine receive the ID of their closest Dijkstra cluster. Here a user given distance threshold has to be passed to reject outlier points with a too large distance to the plot.

### 2.5 QSM reconstruction

#### 2.5.1 QSM basic reconstruction

The SphereFollowing method [4,7,18] utilizes search spheres and onto their surfaces circles are fit in a recursive manner to detect topological ordered cylinders. For details please refer to S1 Text, where a multi paged description of the algorithm exists. One improvement to the previously published SimpleTree implementation is the multitude of available cylinder fitting methods: Center of Mass (COM) with median distance radius [4], Gauss Newton Least Squares (GNLS) [4], Random Sample Consensus (RANSAC) [30], M-estimator SAmple Consensus (MSAC), Randomized M-estimator SAmple Consensus (RMSAC), Maximum Likelihood Estimator SAmple Consensus (MLESAC) [29], Progressive Sample Consensus (PROSAC) [31], Randomized RAndom SAmple Consensus (RRANSAC) [32] and Least MEDian of Squares (LMEDS).

We use the COM method by default as it performs very strong in earlier works [4]. Nevertheless we provide for interested users the GNLS as well as all sample consensus fitting methods provided in the PCL.

Three SphereFollowing main parameters, see user guide for additional information, are predicted and then internally optimized with an automated parameter search [7] utilizing the the downhill simplex method [33, 34]. This parameter search enables researchers to perform the modeling without algorithmic knowledge.

For the parameter search the computer has to decide for two different models which one is better. The best model is chosen via the cloud to model distance. We provide five different distance metrics. The root mean squared distance (RMSD) of each point to the model and the more robust [33] mean distance (MD) of each point to the model. Both RMSD and MD can be used in their more robust form where the distances are cropped to a max value, following the MSAC principle. Lastly the most robust metric out of all is to count the inlier points with less than a threshold distance to the QSM. Good quality clouds should be evaluated with RMSD, those of worse quality with MD. MSAC and the fifth method can be useful if leaf noise exists.

#### 2.5.2 QSM based clustering

We define recursively similar to the *GrowthVolume* from previous works [7]:

**The *GrowthLength (GL)* of a cylinder is the cylinder’s length plus its children’s *GL*.**

We can use the derived QSM after computation to transfer the *GL* of a cylinder to its closest points. Then points can be clustered by their *GL* which takes small values for twig regions and large ones for the root area, Fig 3.

**Fig 3.**
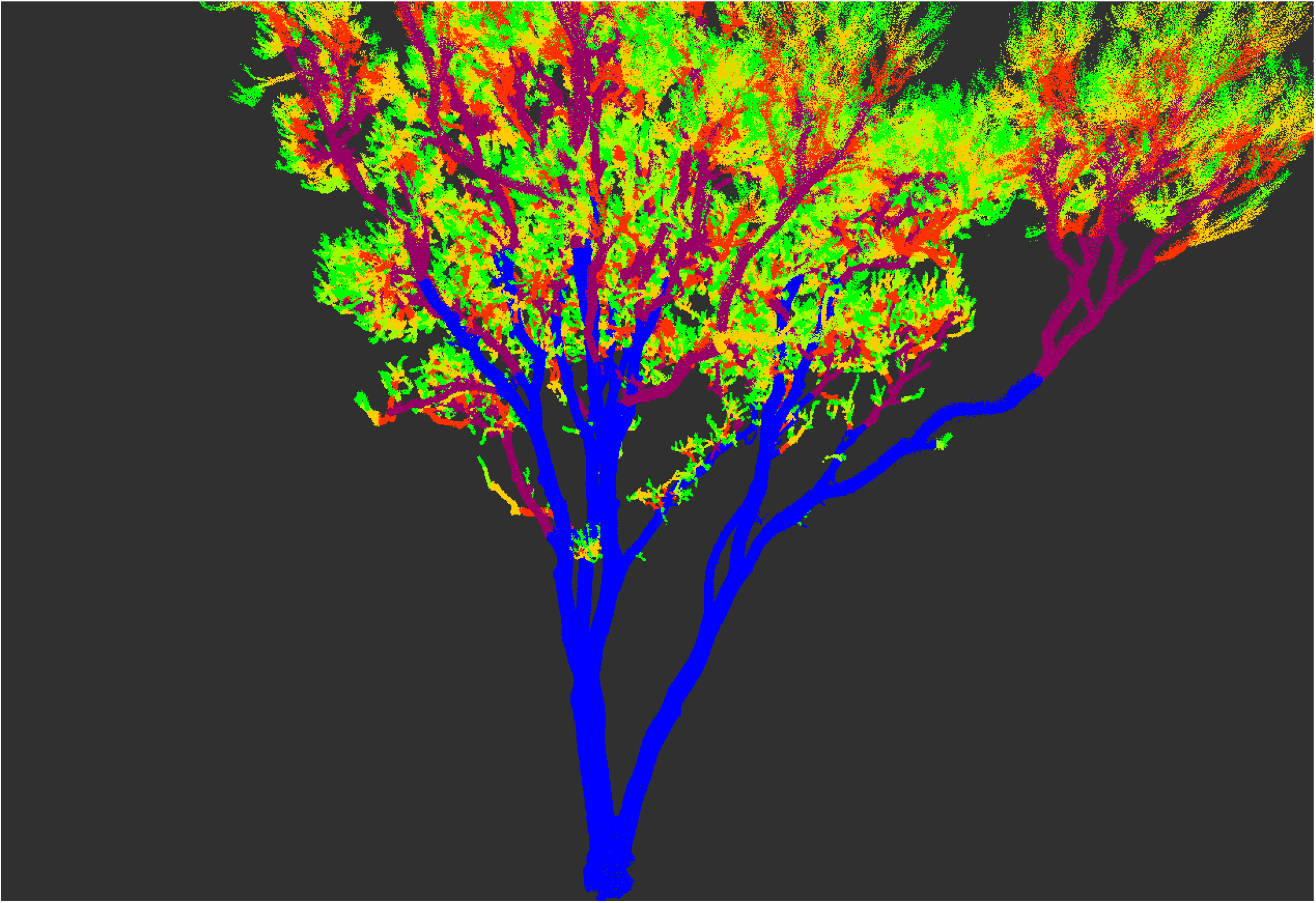
A *GL* clustered point cloud of an *Acer pseudoplatanus*, S4 Dataset. Color codes are ordered from closest to the tips up to close the root: green, orange, red, purple, blue.

#### 2.5.3 QSM advanced reconstruction

In a second modeling run we can use the clusters described in 2.5.2. The three parameters mentioned in 2.5.1 can be estimated and optimized per cluster to receive more accurate cylinders. For example with five clusters we do optimization in a 3 * 5=15 dimensional parameter space.

#### 2.5.4 Detect outlier cylinders

A step which classifies cylinders into WELLFITTED and BADFITTED cylinders exists. Cylinders close to the tips are classified as BADFITTED cylinders [35]. Additionally for the remaining cylinders we compute the average distance of allocated points to the cylinder. Only cylinders with a small average distance are finally classified as WELLFITTED cylinders.

If we look at the average point distance of QSM cylinders we can see that most errors occur either at the twigs or at branch junctions, Fig 4 and Fig 5.

**Fig 4.**
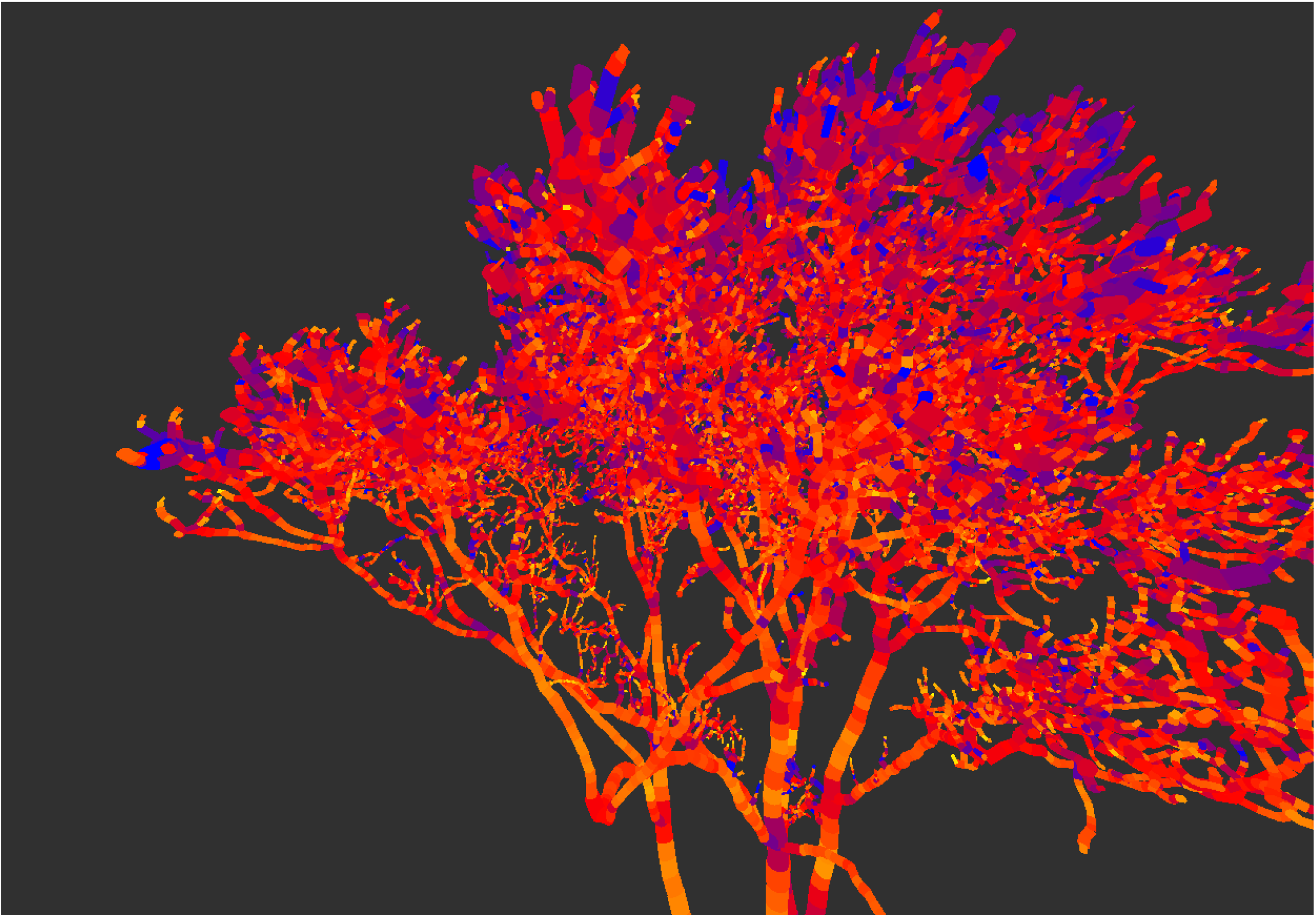
An unfiltered QSM of an *Acer pseudoplatanus*, S4 Dataset. The fitting error of points to their closest cylinder is colorized. Cylinders with low error are colorized orange and cylinders with large errors purple.

**Fig 5.**
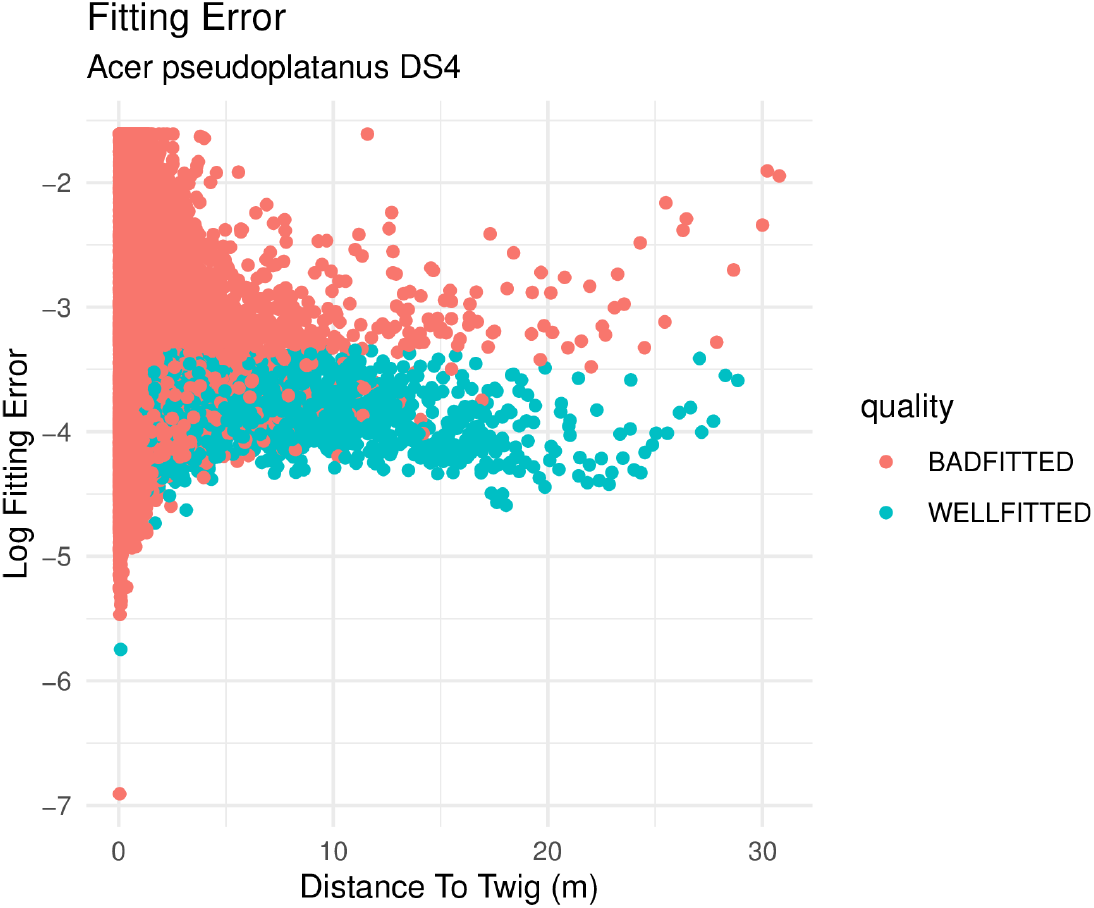
Error plot for an unfiltered QSM of an *Acer pseudoplatanus*, S4 Dataset. The log of fitting error in dependency of the distance to twig.

Some overfitted cylinders can be *Radius* corrected with a QSM median filter, as proposed in earlier works [7]. But even with a good fit quality twig cylinders tend to overestimate the *Radius* [35]. Footprint size of the laser and co-registration errors lead to the fact the cloud regions of small diameter components tend to overestimate.

### 2.6 QSM filters

#### 2.6.1 *Tapering* filter

We implemented a taper correction based on the one TreeQSM [42, 43] provides.

#### 2.6.2 *GrowthVolume* (GV) filter

We applied a *GV* filter in [7] which handled the overestimation to some extend. There we predicted the *Radius* from the *GV* with the Eq 1.

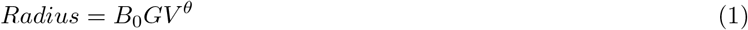

#### 2.6.3 *VesselVolume* (VV) filter

The *GV* of a cylinder is always effected by the error of twigs as the supported twig *Volume* is included in its value. By it’s definition the *GV* is therefore not the best predictor. We can be more robust against twig *Radius* overestimation by ignoring the *Radii* with the following definitions:

**We define the *Reverse Pipe Area Branchorder (RPAB):* The WBE model considers that the branching architecture is self-similar [16]. In a self-similar branching architecture the *Radius* as well as the cross sectional area of a twig is constant. As we do not know this area in** cm^2^ **we define that *RPAB* is a tree specific area unit which equals one for a tip cylinder. By following the pipe model theory [14] the *RPAB* of a cylinder at any location inside the QSM equals the number of supported tips. Supported tip cylinders always have a recursively defined parent relation to the query cylinder**.

The *RPAB* serves as a proxy for the cross sectional area but is by the definition independent from any *Radius* measurement.

To retrieve a *Radius* independent proxy for the *Volume*, we define:

**The *VV* of a cylinder is the cylinder’s *RPAB* multiplied with the *Length* plus its children’s *VV*.**

We can see in Fig 6 that the VV is invariant during the correction and GV is not. Also notable is that outliers are less strong visible for VV.

**Fig 6.**
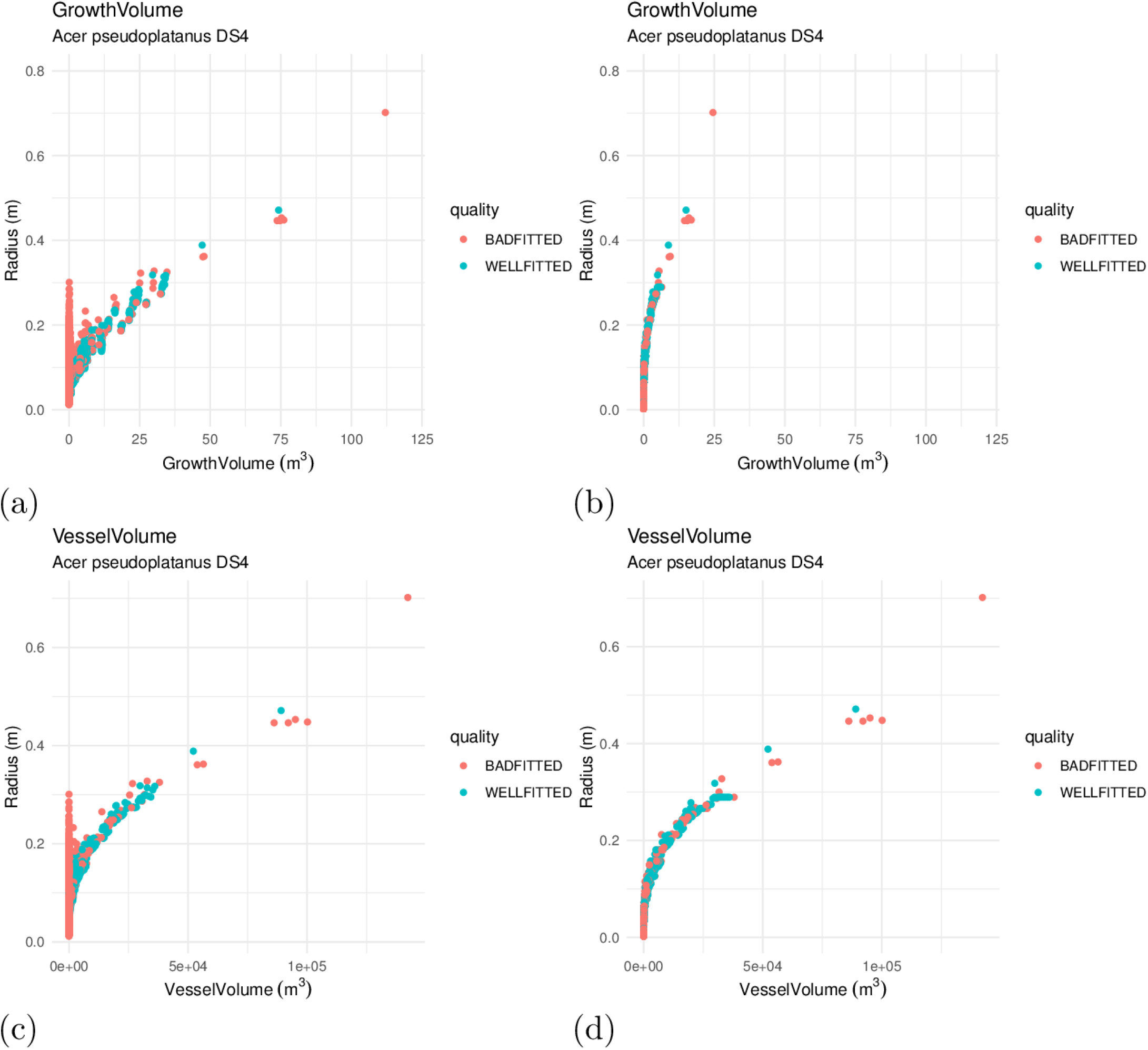
Allometric plots of an *Acer pseudoplatanus*, S4 Dataset. (a) The *GV* extracted from an uncorrected QSM. (b) The *GV* from the same QSM which has been corrected. The magnitude of the predictor was effected by the correction. (c) The VV of the uncorrected QSM. (d) The VV of the corrected QSM covers the same value range.

SimpleForest provides such a *VV* predicted *Radius* correction utilizing Eq 2, with intercept *B*_1_ estimation being optional.

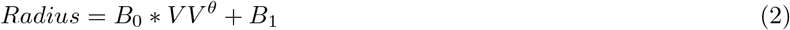

The additive component *B*_1_ is relevant in case twig regions have been removed during denoising.

#### 2.6.4 *Reverse Pipe Radius Branchorder* (RPRB) filter

Doing the non linear underlying power function fit unsupervised can lead to the termination in a non desired local minimum if the initial values for *B*_0_ and *θ* are not good enough. We get an easier mathematical model by using a *Radius* independent proxy for the *Radius* itself:

**The *RPRB* is the square root of the *RPAB*.**

This relation is a linear model of a simple form as also the intercept is forced to the origin here:

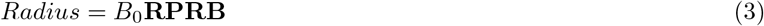

Exemplary data into which equation 3 is fitted can be seen in Fig 7.

**Fig 7.**
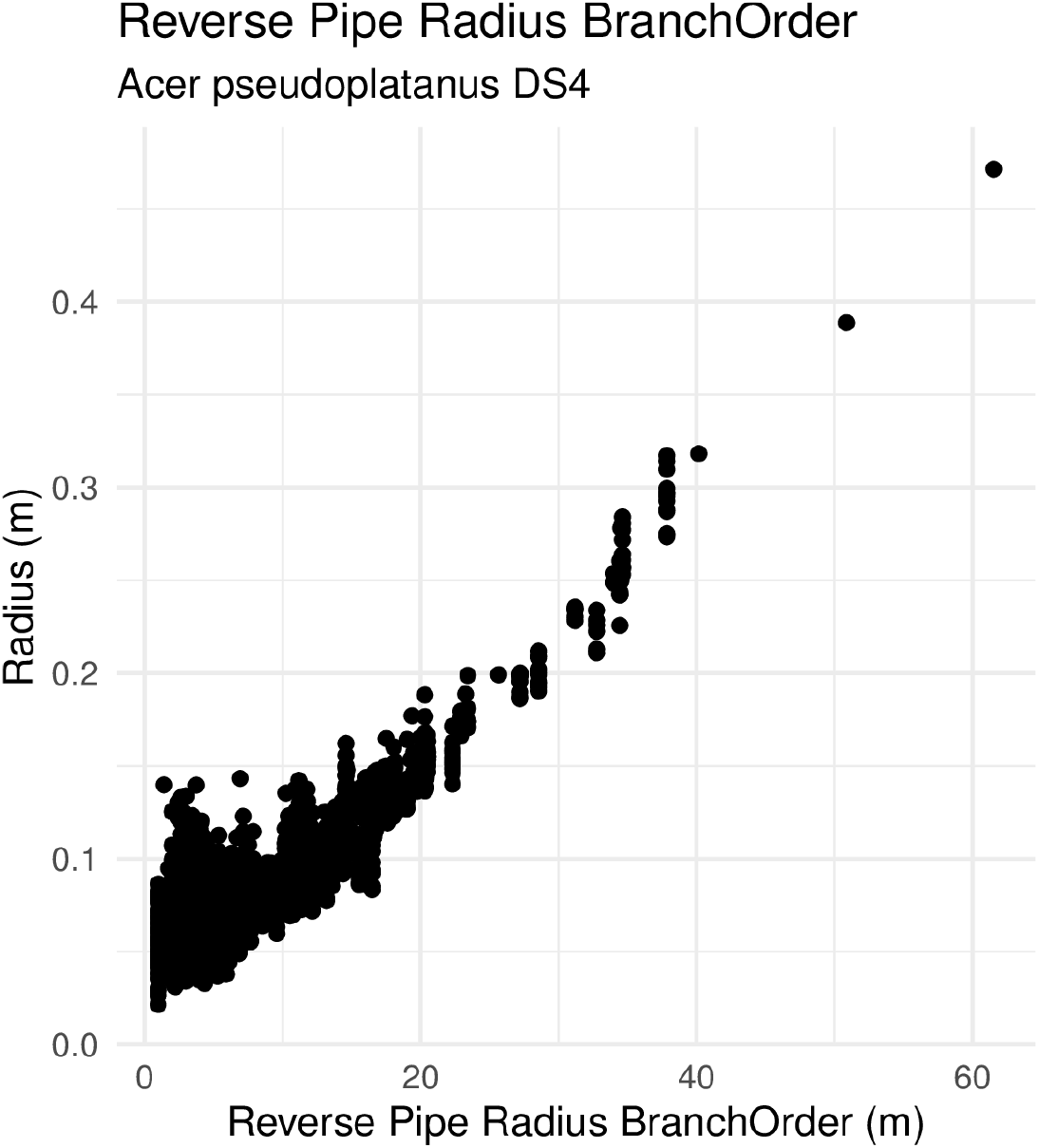
RPRB Scatter plot for an *Acer pseudoplatanus*, S4 Dataset. Plot of the *RPRB* on the x-axis and *Radius* as a predicted value.

#### 2.6.5 Refitting cylinders

As we only have fitted circles with the SphereFollowing method, we can align points to their closest cylinder and do a cylindrical refit on those points [4], Fig 8.

**Fig 8.**
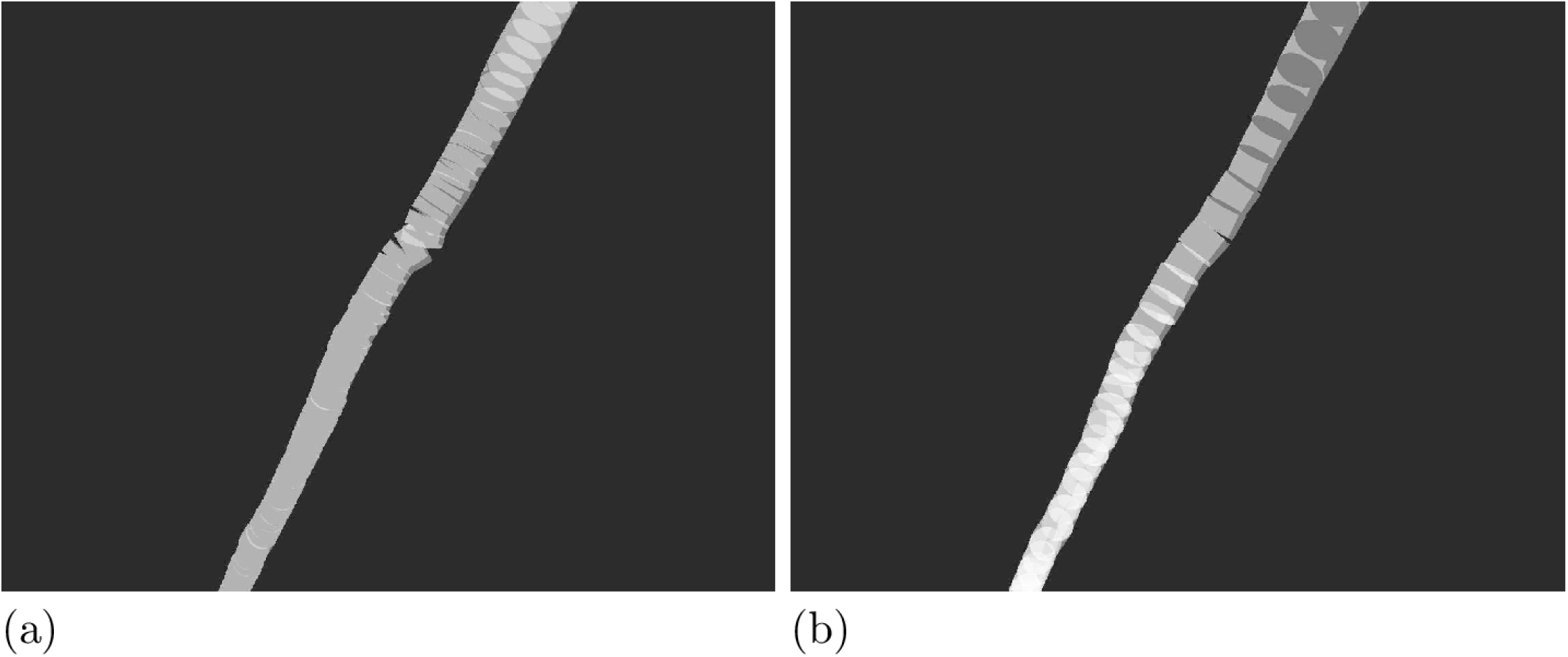
Cylinder correction for an *Prunus avium*, S1 Dataset. (a) After the SphereFollowing method the cylinders can still have larger errors. (b) The cylinder refit step was performed on the same QSM and the before visible error is gone.

## 3 Results

The *Quercus petraea* and denoised *Pinus massioniana* of S1 Dataset were corrected with the *Tapering* and *Erythrophleum fordii* were corrected with *VV* approach. In a second modeling run all QSMs were corrected with the *RPRB* filter. For S2 Dataset [36] we used the *VV* filter in one run and the *RPRB* filter in a second one. For S3 Dataset we applied *Tapering* firstly and then in a second run the *RPRB* filter, Fig 9.

**Fig 9.**
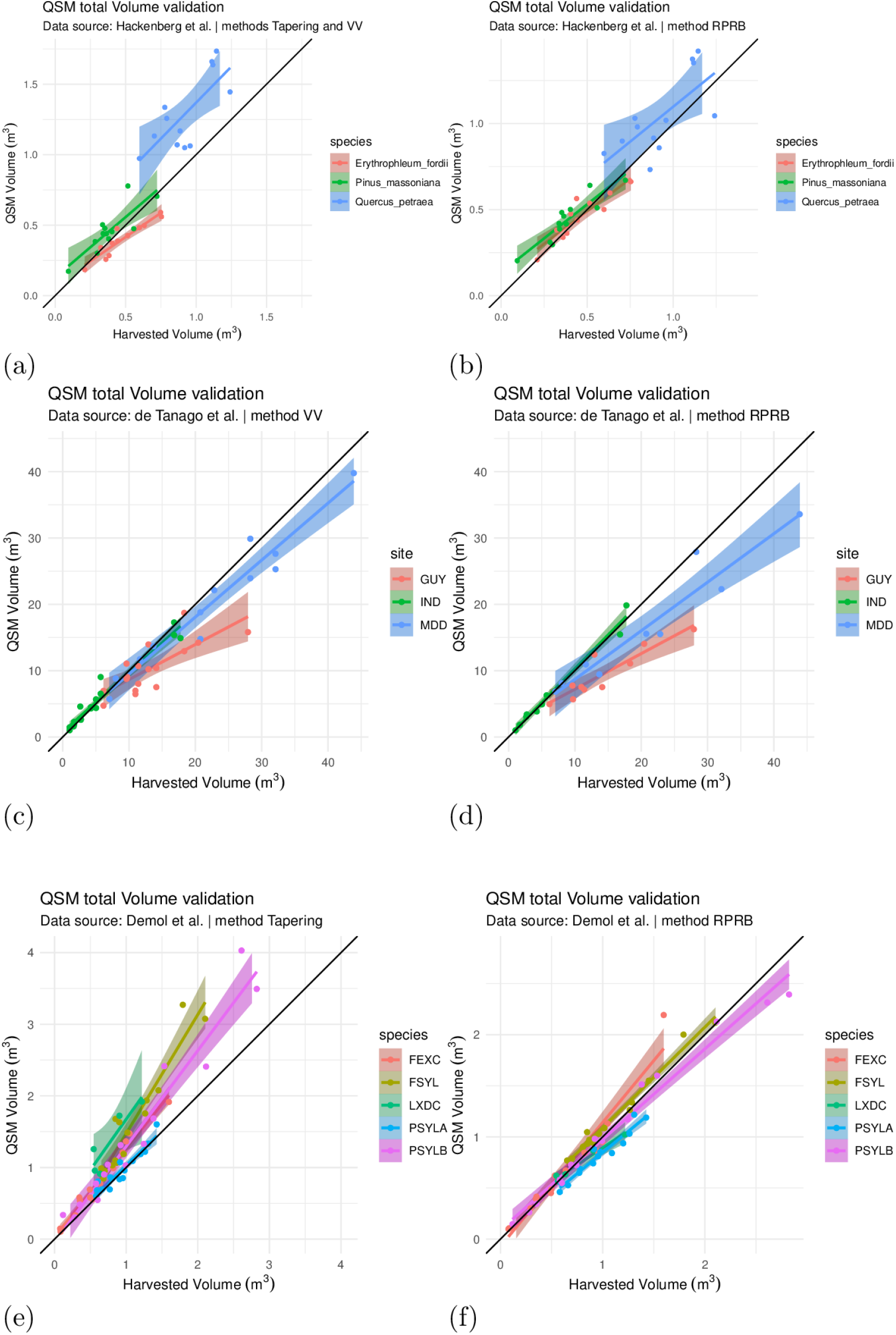
Volume validation for S1 Dataset, S2 Dataset and S3 Dataset. Modeling results of (a) S1 Dataset with *Tapering* and *VV* correction, (b) S1 Dataset with *RPRB* correction, (c) S2 Dataset with *Tapering* correction, (d) S2 Dataset with *RPRB* correction, (e) S3 Dataset with *VV* correction, (f) S3 Dataset with *RPRB* correction.

We show for S3 Dataset within one plot SimpleForest results correct one time with *Tapering* and one time with *RPRB* compared to TreeQsm [42] and in a second plot SimpleForest *Tapering* results which have been filtered by using both tools. SimpleForest *Tapering* corrected results have to be in a 5% range to TreeQSM *Tapering* corrected results, otherwise the QSM is filtered out, Fig 10.

**Fig 10.**
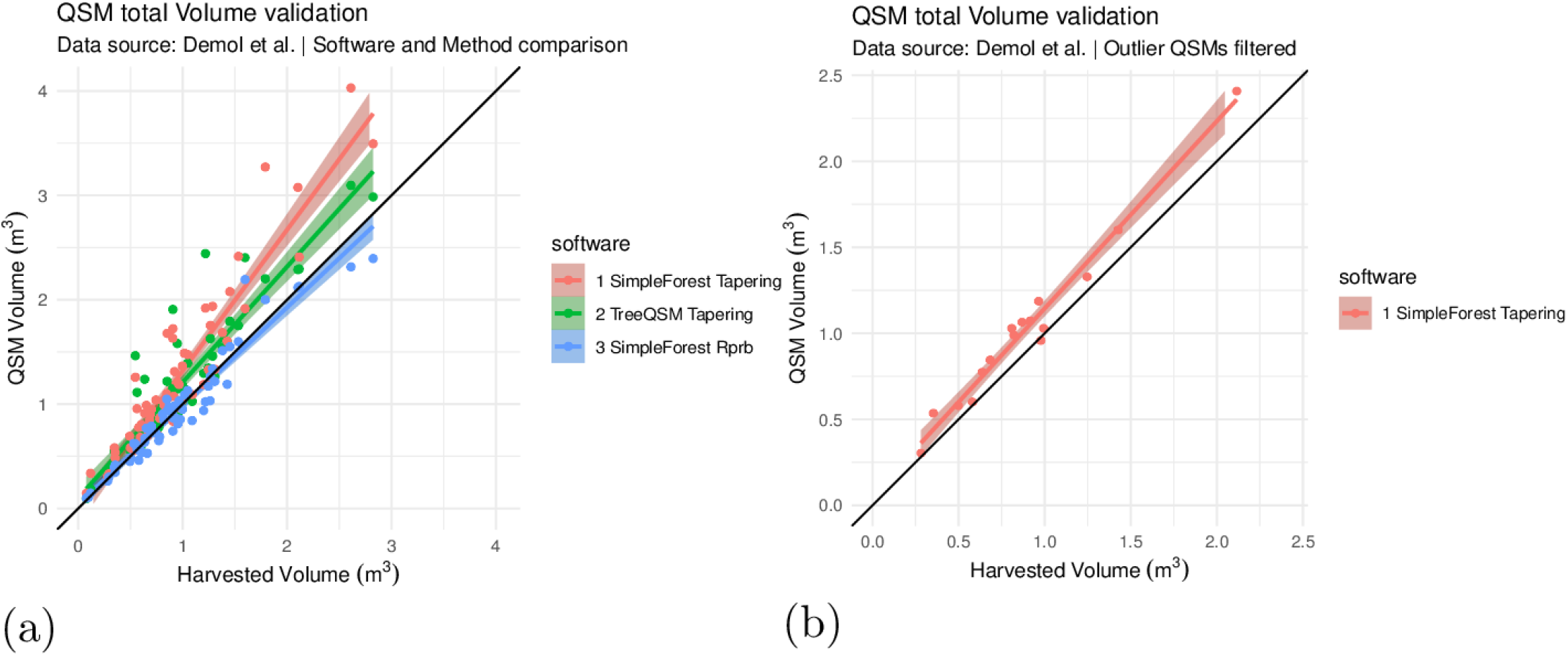
Software and method comparison, S3 Dataset. (a) Comparison of SimpleForest *Tapering*, TreeQSM *Tapering* and SimpleForest *RPRB* results for S3 Dataset. (b) We only see SimpleForest *Tapering* volume predictions which are close to TreeQSM *Tapering* volume prediction, other QSMs are filtered out.

Combining SimpleForest results for all data sets with the *VV* approach for S2 Dataset and *RPRB* filter otherwise leads to Fig 11.

**Fig 11.**
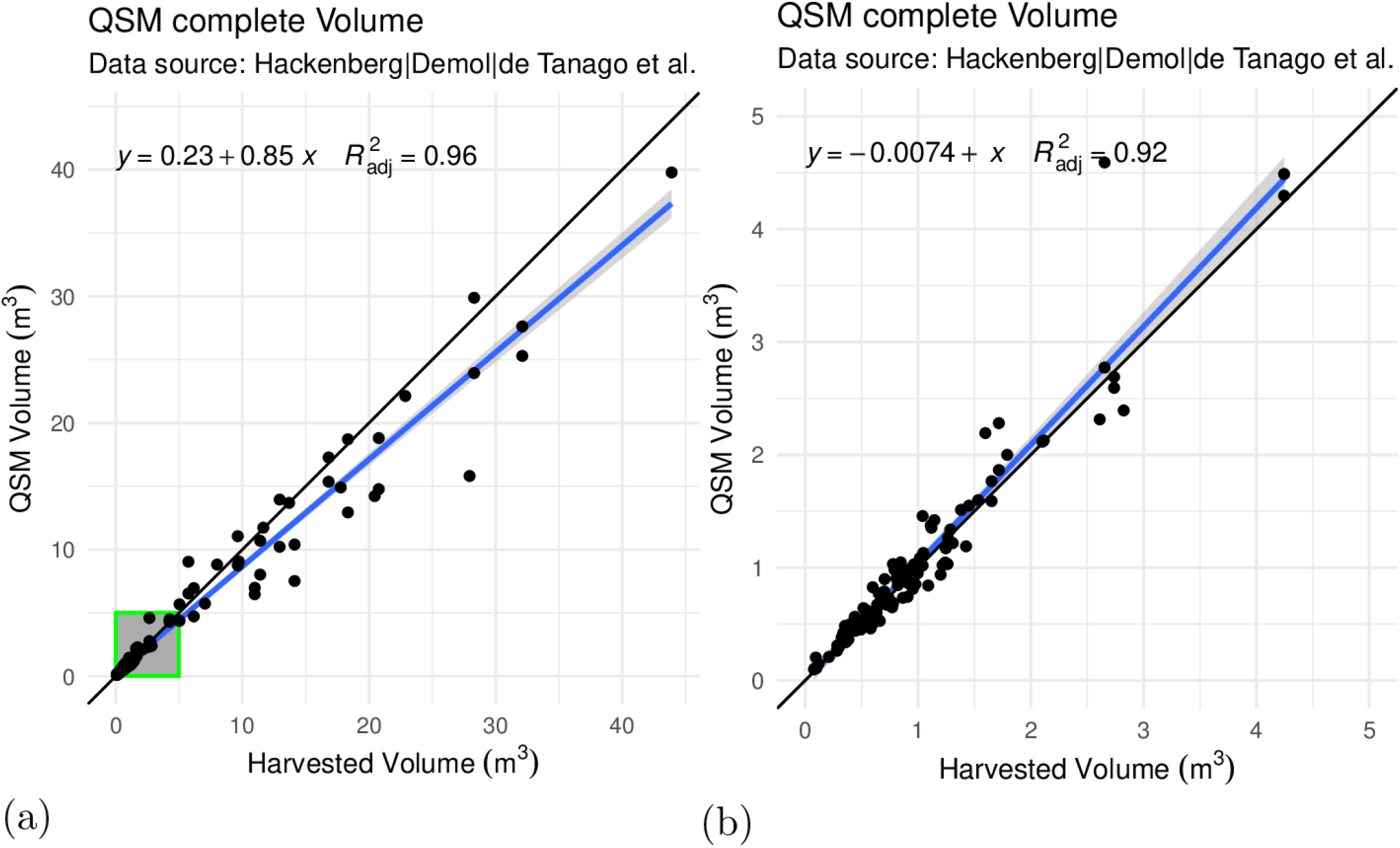
Combined S1 Dataset, S2 Dataset and S3 Dataset validation. (a) Combined evaluation of all three data sets showing a zoom area of interest used for the next figure. (b) The zoom view showing trees with a maximum volume of 5 *m*^3^.

We provide a summary of the results in table 1.

**Table 1.** Numerical validation of QSM volume measures compared to harvested reference volume measures.

| Data | Software | $RMSE(m^3)$ | $RMSE_{rel.}$ | $CCC$ | $r^2_{adj.}$ |
| --- | --- | --- | --- | --- | --- |
| S1 Dataset [35] | SimpleForest <sup>a</sup> | 0.12 | 0.10 | 0.91 | 0.87 |
| S1 Dataset [35] | SimpleForest <sup>b</sup> | 0.24 | 0.16 | 0.79 | 0.83 |
| S2 Dataset [36] | SimpleForest <sup>c</sup> | 3.25 | 0.08 | 0.94 | 0.92 |
| S2 Dataset [36] | SimpleForest <sup>a</sup> | 4.67 | 0.14 | 0.87 | 0.90 |
| S3 Dataset [37] | SimpleForest <sup>a</sup> | 0.14 | 0.06 | 0.97 | 0.93 |
| S3 Dataset [37] | SimpleForest <sup>d</sup> | 0.43 | 0.11 | 0.80 | 0.88 |
| S3 Dataset [37] | TreeQSM <sup>c</sup> | 0.32 | 0.11 | 0.86 | 0.85 |
| S3 Dataset [37] | SimpleForest <sup>e</sup> | 0.15 | 0.08 | 0.94 | 0.98 |
| S1 Dataset, S2 Dataset, S3 Dataset [35–37] | SimpleForest <sup>f</sup> | 1.85 | 0.05 | 0.97 | 0.96 |

<sup>a</sup> = *RPRB* filtering
<sup>b</sup> = *VV* filter for *Erythrophleum fordii*, *Tapering* filter otherwise
<sup>c</sup> = only *VV* filter.
<sup>d</sup> = only *Tapering* filter.
<sup>e</sup> = only *Tapering* filter, rejected QSMs not in a 5 % range to TreeQSMs.
<sup>f</sup> = *VV* filter for S2 Dataset data, otherwise *RPRB* filter.

Our plot data did not have ground truth attached, but we use it for visually inspecting the quality of the DTM, Fig 12.

**Fig 12.**
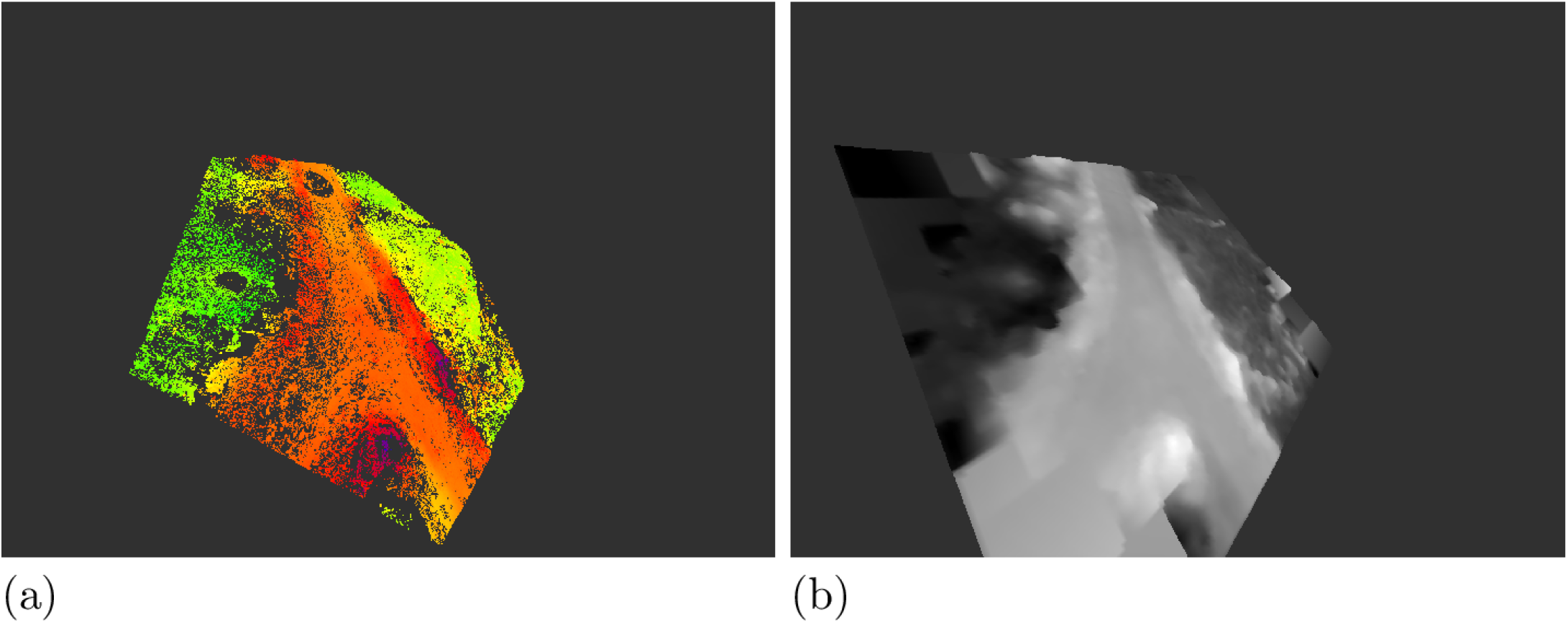
Visual DTM quality assessment, S5 Dataset. (a) We see the height of ground points visualized increasingly from green to red. A forest road is visible. (b) The DTM height values are colored increasingly from dark to bright. The DTM captures the road without artifacts.

The modeled QSMs also do not show segmentation errors in Fig 13.

**Fig 13.**
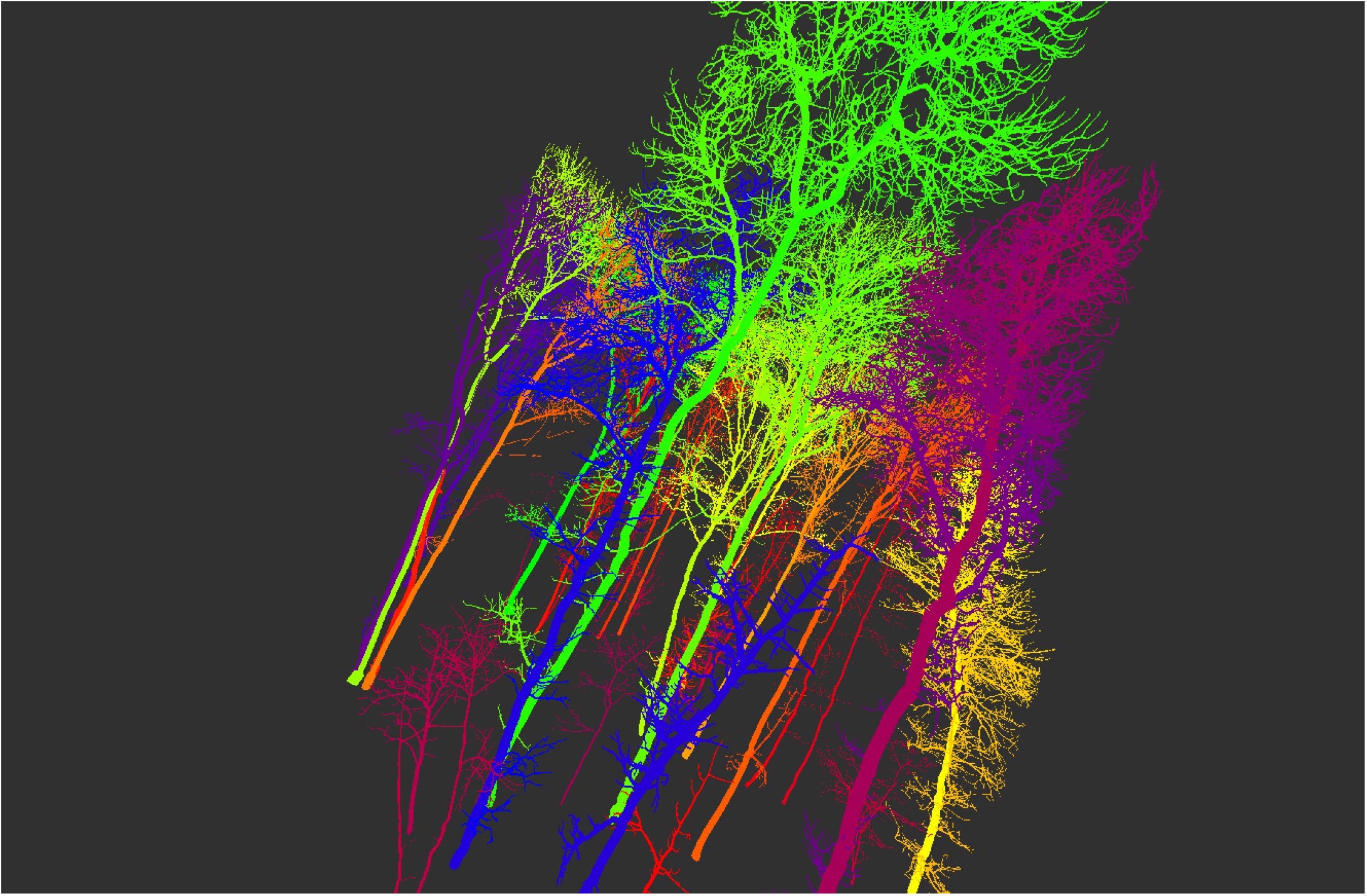
Final QSMs of S5 Dataset. Each QSM is colored differently.

The *Acer pseudoplatanus* QSM, S4 Dataset, contains more than 35k cylinders. We use this QSM to show some statistical analysis potential in Fig 14.

**Fig 14.**
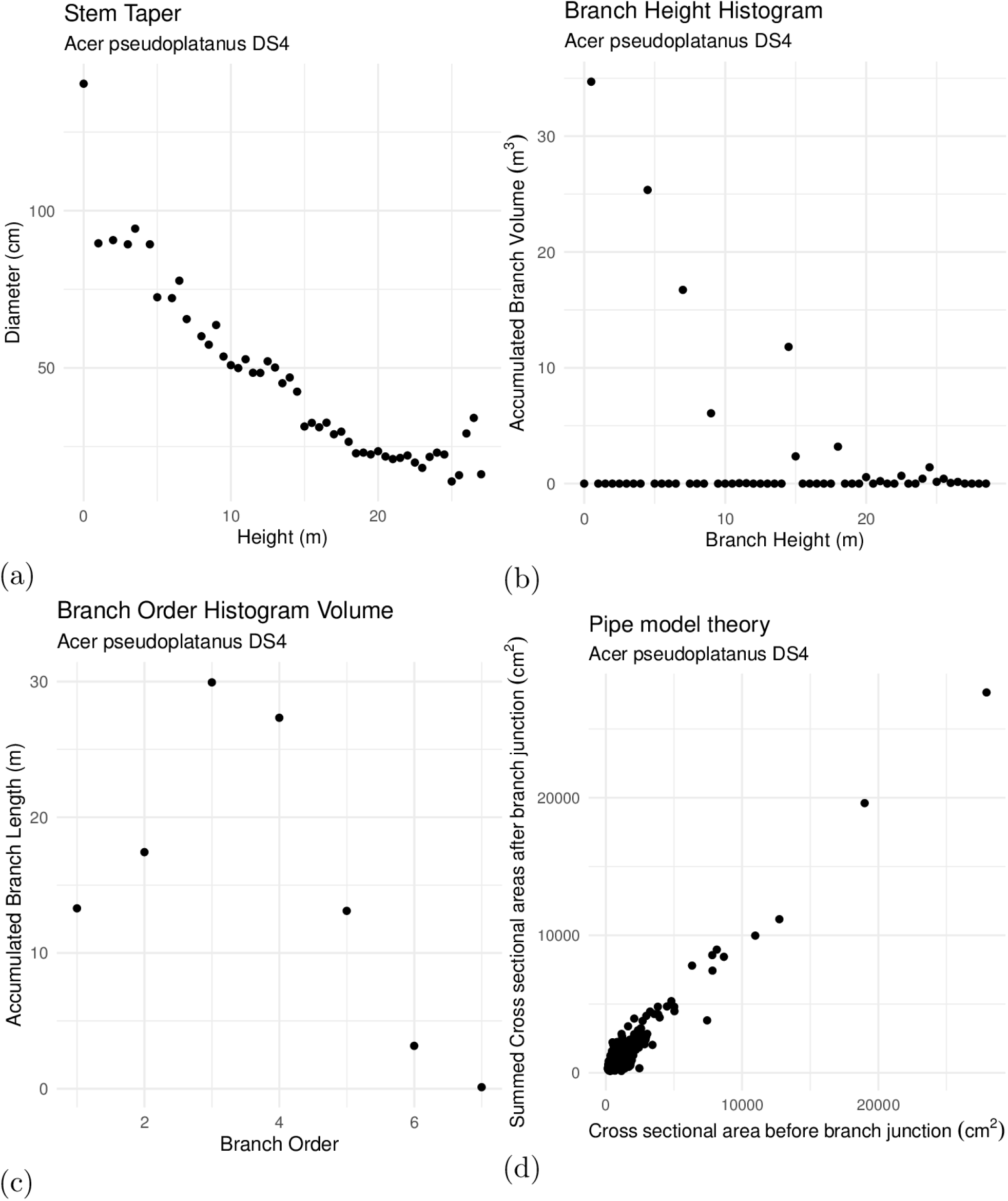
Exemplary statistic plots for an *Acer pseudoplatanus*, S4 Dataset. (a) We show a traditional stem taper. (b) The branch volume distribution binned along the height of the branch growing out of the stem. The maximum can define the crown base. (c) The branch volume accumulated for each branch order. (d) An analysis in tradition of the Pipe Model Theory [7].

## 4 Discussion

SimpleForest utilizes an advanced parameter search for optimizing the SphereFollowing method compared to the brute force search published in earlier works [7] leading to better models. Nevertheless, as input data already overestimates the twig regions [35] a radius correction has to be applied. The choice of this correction influences the results significantly. On the S3 Dataset data set we can perform a direct comparison versus an earlier published approach and our *RPRB* filter performs very strong as can be seen in Fig 10 (a).

We can also see, that even with the less well performing *Tapering* radius correction we can achieve better results, if we use more than one QSM tool. Outlier QSMs can be simply filtered out if the QSMs of different tools do not have similar volume predictions, Fig 10 (b). In this work we only use TreeQSM, but also 3D Forest [40] can build QSMs nowadays. By using all three tools a more sense of uncertainty can be gained and filtering out QSMs from tree clouds which are hard to model can be automated.

The S1 Dataset was the second data set revealing the strength of the RPRB filter. Only for S2 Dataset this filter performed worse than the comparison filter. The reason behind this performance is simply, that we lost the thin branching structure during the de-noising routine, but forced intermediate sized branches to become thin, Fig 15.

**Fig 15.**
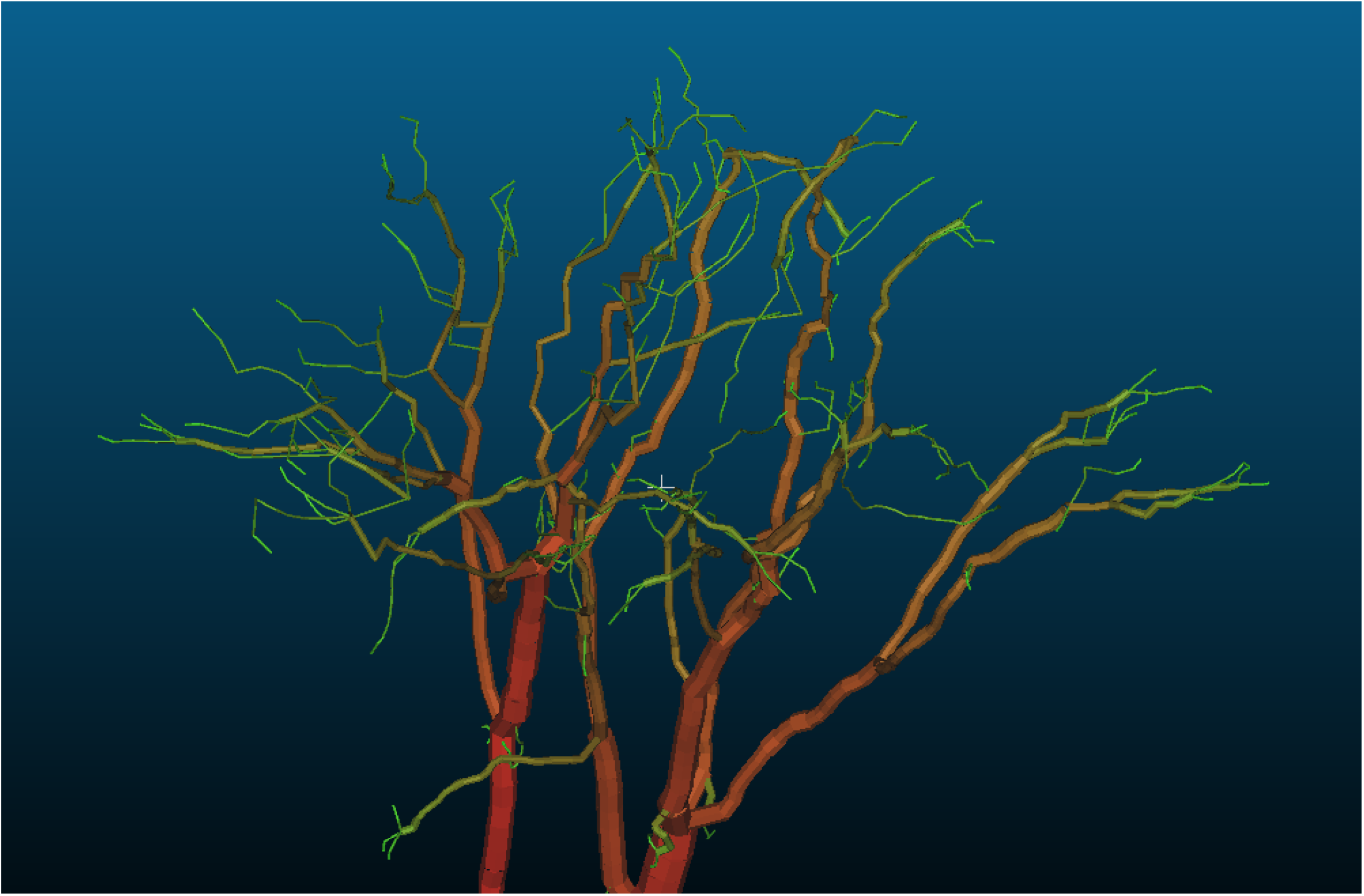
Underestimation of twigs, S2 Dataset. Intermediate branches have been underestimated after RPRB correction for two S2 Dataset trees, QSMs visualized with Cloud Compare. The tree clouds lost the twig points during denoising.

As non TLS derived data to evaluate DTM and segmentation quality was not available, we have to rely on visual inspection. Our example plot reveals no errors and we do not have to improve with manual interaction our segmentation results [41].

Our QSM validation results are in line with other available tools and sometimes even stronger than results which have been published by those. At least for leaf-off conditions the complete data processing can be fully automatized on a plot level. If leaf-on conditions are met, we provide two functional semi automatic leaf filtering procedures.

Regarding that the embedding platform Computree has other plugins for forest modeling in addition and that we provide as much interfaces as possible to external software tools, we recommend using this software suite for estimating tree’s above volume as well as gaining insight into forest ecology relevant questions.

## 5 Availability and Future Directions

### 5.1 Future work

The *RPRB* filter should not rely on an origin forced intercept, as twigs can be removed during leaf separation. Another downside of this filter is the fact, that the *RPRB* is same for all cylinders inside a segment [7]. Notice, that we defined a segment in previous works as the list of all cylinders between two neighboring branch junctions. The *VV* filter does not have this downside and if the non linear fit can be performed in a more robust way even better filtering might be doable.

Lastly we consider it beneficial to gather with our semi automatic leaf separation approaches training data for a full automatic leaf separation procedure. Tiny dnn [44, 45] is a deep learning header only library which allows easy integration into SimpleForest code base. A network architecture using FFPHs as input can be implemented fastly.

## Acknowledgments

S5 Dataset is provided by the Leipzig Canopy Crane facility financed by the German Centre for Integrative Biodiversity Research (iDiv) Halle-Jena-Leipzig and the instute of Systematic Botany and Functional Biodiversity, Institute for Biology, Leipzig University.

## Author Contributions

Hackenberg Jan: Conceptualization, Data Curation, Formal Analysis, Investigation, Methodology, Resources, Software, Validation, Visualization, Writing – Original Draft Preparation, Writing – Review & Editing

Calders Kim: Data Curation, Writing – Review & Editing

Miro Demol: Data Curation, Validation, Writing – Review & Editing

Pasi Raumonen: Methodology, Writing – Review & Editing

Alexandre Piboule: Software, Resources, Writing – Review & Editing

Disney Mathias: Conceptualization, Data Curation, Writing – Review & Editing

